# FastFeatGen: Faster parallel feature extraction from genome sequences and efficient prediction of DNA *N*^6^-methyladenine sites

**DOI:** 10.1101/846311

**Authors:** Md. Khaledur Rahman

## Abstract

*N*^6^-methyladenine is widely found in both prokaryotes and eukaryotes. It is responsible for many biological processes including prokaryotic defense system and human diseases. So, it is important to know its correct location in genome which may play a significant role in different biological functions. Few computational tools exist to serve this purpose but they are computationally expensive and still there is scope to improve accuracy. An informative feature extraction pipeline from genome sequences is the heart of these tools as well as for many other bioinformatics tools. But it becomes reasonably expensive for sequential approaches when the size of data is large. Hence, a scalable parallel approach is highly desirable. In this paper, we have developed a new tool, called `FastFeatGen`, emphasizing both developing a parallel feature extraction technique and improving accuracy using machine learning methods. We have implemented our feature extraction approach using shared memory parallelism which achieves around 10× speed over the sequential one. Then we have employed an exploratory feature selection technique which helps to find more relevant features that can be fed to machine learning methods. We have employed Extra-Tree Classifier (ETC) in `FastFeatGen` and performed experiments on rice and mouse genomes. Our experimental results achieve accuracy of 85.57% and 96.64%, respectively, which are better or competitive to current state-of-the-art methods. Our shared memory based tool can also serve queries much faster than sequential technique. All source codes and datasets are available at https://github.com/khaled-rahman/FastFeatGen.

## 1 Introduction

*N*^6^-methyladenine (6mA) is very common in prokaryotes whose primary functions lie in the host defence system [20]. It is an abundant modification in mRNA which has also been found in many multicellular eukaryotes such as *Caenorhabditis elegans* [13] and *Drosophila melanogaster* [32], and hence it has been proposed as a new epigenetic marker in eukaryotes [20]. Some studies have revealed that 6mA can control the acuity of infection and replication of RNA viruses like HIV and Zika virus [16, 17]. A recent study demonstrates that 6mA modification can be heavily present in human genome and depletion of 6mA may lead to tumorigenesis [30]. Identification of 6mA in the genome will be helpful to characterize many biological functions and drug discovery.

Several experimental approaches exist to identify 6mA in genome. As described in [20], an antibody against *N*^6^-methyladenine can identify *N*^6^-adenine methylation in eukaryotic mRNAs which can further be used to identify *N*^6^-methyladenine in DNA [11]. This technique is ambiguous due to the fact that other adenine-based modifications can be recognized. Liquid chromatograpy coupled with tandem mass spectrometry gives another comprehensive approach to identify 6mA [10]. Some restriction enzymes are sensitive to DNA methylation to differentiate between methylated nucleotides and unmethylated nucleotides which can be used to identify 6mA [25]. Single Molecule Real Time (SMRT) sequencing can determine kinetics of nucleotides during synthesis [9]. It has been applied to map 6mA and 5mC at the same time in *Escherichia coli* [7]. Noticeably, it can not differentiate between 6mA and 1mA, though this technique is very expensive. There are other experimental methods in the literature which have been found effective, e.g., liquid chromatography coupled with tandem mass spectrometry [13], and capillary electrophoresis alongside laser-induced fluorescence (CE & LIF) [14].

Most of the existing experimental methods are time consuming and expensive as mentioned above. Since the distribution of 6mA sites in the genome is not random and can follow some patterns, computational methods may be efficient and cost-effective. There are few such methods (6mA-Pred [4] and iDNA6mA-PseKNC [8]) which help to identify 6mA sites using supervised machine learning approaches. But, these methods adopt a sequential approach to extract features from DNA sequences which often slow down the process. Recently, convolutional neural networks (CNN) model has been applied to this problem [29]. However, comparison is not fair or ambiguous as jackknife testing is performed in 6mA-Pred whereas Tahir et al. use 20% samples of the dataset. Hence, we exclude this method from comparison due to inconsistency. There is still a need for a robust and precise tool that can facilitate faster and efficient identification of 6mA sites.

Various tools exist that extract features from DNA/protein sequences for prediction purposes, e.g., some tools extract features from genomic sequences to predict on-target activity in CRISPR/Cas9 technology [6, 22] whereas other tools extract features from protein sequences to make efficient predictions [5, 21, 23, 24]. But, almost all authors use a sequential approach [2, 18, 19] for feature extraction. With the advent of multicore processors [1, 26], a single machine can have two or more processing units which can lead to a significant speed-up of a properly written program. In this paper, we introduce such a parallelization technique in `FastFeatGen` (**Fast**er **Feat**ure extraction from **Gen**ome sequences) to extract features from DNA sequences which can also be applied to RNA/protein sequences as well with small modification. So, feature extraction techniques from large scale datasets will be significantly accelerated by our tool.

Over the years, a plethora of supervised machine learning based methods have been applied to solve several bioinformatics problems [15, 28]. However, to the best of our knowledge, Chen et al. [4] were the first to tackle identification of 6mA sites in rice genome using Support Vector Machine (SVM). In this paper, we advance this concept with faster feature extraction and selection techniques and several supervised learning methods to achieve better performance. We also apply widely used neural network models to this problem in both supervised and unsupervised ways. Our extensive experimental results show that unsupervised CNN model is unable to surpass supervised models. This is likely due to the small size of the datasets, and, more interestingly, Extra-Tree Classifier (ETC) proposed by [12] performs very well despite its small set of features. We summarized our contributions as follows:

- We have introduced faster approaches for feature extraction techniques from genome sequences (Section 2.2).
- We have applied a lucid feature selection technique to select important and relevant features (Section 2.4).
- We have performed an extensive set of experiments using both supervised and unsupervised machine learning methods to find a robust model (Section 2.3).
- We have compared our results with other state-of-the-art methods to show effectiveness and efficiency of our model (Section 3).
- We have also analyzed the processing time of new query sequences (Section 3.5).

## 2 Methods

The workflow of our tool is shown in Fig. 1. At first, features are extracted from input genome sequences (datasets). Then, relevant features are selected using the appropriate technique. After that, selected features are fed to supervised machine learning methods to build the predictor. At this stage, several parameters are tuned until the model is optimized. Many existing methods follow Chou’s 5-step rules (see [5]), and our tool is analogous to it. We describe each of these steps below.

**Fig. 1:**
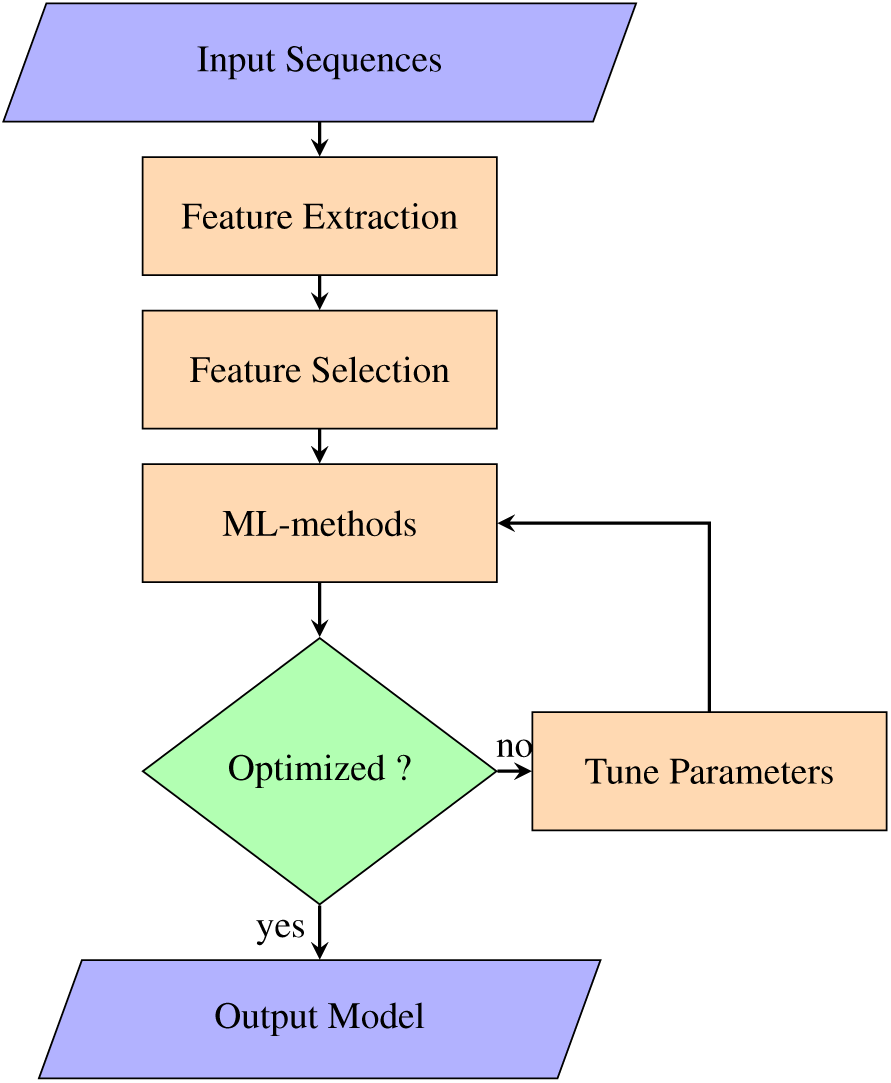
Flowchart for `FastFeatGen`. Input Sequences - dataset containing DNA sequences, Feature Extraction - extract features from DNA sequences, Feature Selection - select relevant features using feature importance score, ML-methods - apply supervised machine learning methods on selected features, Optimized - check whether model is better or not, Tune Parameters - tune several parameters in ML-methods like learning rate, kernel function, etc., Output Model - produce optimized model for prediction.

### 2.1 Datasets

We have obtained two balanced datasets, Dataset1 and Dataset2 from [33] and [31], respectively. Dataset1 contains 880 positive samples (6mA sites) and 880 negative samples (non-6mA sites). Positive samples are from the rice genome, which are available at NCBI Gene Expression Omnibus^1^ with the accession number GSE103145. Dataset2 contains 1934 positive samples and 1934 negative samples. Positive samples are curated from *Mus musculus* genome which are available in MethSMRT database [31]. Each sequence of both datasets is 41-bp long and nucleotide “A” is present at the center. More details about these datasets and negative samples generations can be found in [4] and [8].

We represent a 41-bp sequence by *S* = *s*_1_*s*_2_*s*_3_ … *s*_41_, where *s*_*i*_ represents a nucleic acid in sequence *S* and 1 ≤ *i* ≤ 41. Thus a dataset is represented by *D* = *S*_1_*S*_2_*S*_3_ … *S*_|*D*|_, where |*D*| is the size of dataset containing both positive and negative samples. For both datasets, *S*_1_ … *S*_|*D*|/2_ are positive samples and *S*_|*D*/2|+1_ … *S*_|*D*|_ are negative samples.

### 2.2 Feature engineering

Features are the heart of supervised machine learning methods. We manually extract different types of information from given datasets and feed that information along with their corresponding true class to machine learning algorithms for training and testing purposes. In this paper, we generate four types of features which are discussed below.

#### Nucleic acids composition (NAC)

This is also known as position independent features or *k*-mers or *n*-grams. Each sequence may have certain short length patterns (also known as motifs) of NACs which are consistent over the whole dataset and so they may contribute to the learning model. In this technique, normalized frequency of a composition of nucleic acids is considered in corresponding sequence and finally a feature vector is constructed for the whole dataset. Length of a composition of nucleic acids is determined by order. For example, if order is 2, then all compositions of two nucleic acids is considered to extract features and a single feature vector is constructed for each composition. We normalize the frequency dividing by length of the sequence. We can define it mathematically as following:

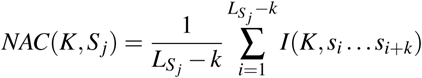

where *K* is a *k*-mer, *S* is a sequence, *L*_*S*_*j*__ represents the length of *S*_*j*_ which is *j*^*th*^ sequence of the whole dataset, and *I*(.) is an identity function which returns 1 when *K* is same as *s*_*i*_ … *s*_*i*+*k*_; otherwise, it returns 0.

### Position specific features (PSF)

Position of a motif in a genome sequence may carry important information which can be found consistent over the whole dataset. Undoubtedly, this position specific information can contribute to the learning model. In this type of features, a binary feature vector is constructed by checking the presence of a *k*-mer in certain position over the whole sequence. We can also define it mathematically as following:

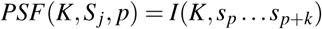

where, *p* is the starting position of *k*-mer and all other variables carry same meaning as NAC.

#### Di-gapped nucleotides (DGN)

Sometimes a gap between relative position of two amino acids in a protein sequence carry important information which may also contribute to supervised learning methods. We are motivated by this technique and use it to extract features from genome sequences as well. We construct a feature vector for each composition of two gapped nucleotides by normalizing its frequency in a sequence. We can formally define DGN as the following:

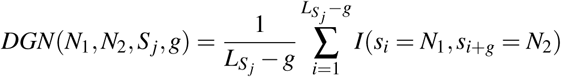

where, *g* is the gap length, *N*_1_ and *N*_2_ are two nucleotides. *I*(.) is an identity function which returns 1 if *i^th^* symbol of sequence *S*_*j*_ is *N*_1_ and (*i* + *g*)^*th*^ symbol of *S*_*j*_ is *N*_2_; otherwise, it returns 0.

#### Bayesian posterior probability (BPP)

In this technique, we first calculate the normalized frequency of each 2-mer for each position over the whole dataset. As we consider 2-mer, there are 40 different positions in a 41-bp sequence and we can have a total of 16 2-mers from all possible combinations of nucleotides. We construct a 40 × 16 matrix for the positive and negative samples separately. Then we extract BPP features in the following way: for each sequence we create a vector of size 80 where first 40 entries represent the posterior probabilities of position specific 2-mer in positive samples and last 40 entries represent the posterior probabilities in negative samples. More details of this approach can be found in [27].

#### Parallelization in feature extraction

We parallelize all the above feature extraction techniques in shared memory parallelism, which is accomplished through Single Instruction Multiple Data (SIMD) computing combined with multithreading. In this approach, instead of one sequence at a time, we pass *nt* sequences at a time to extract features using *nt* cores. Figure 2 shows a schematic diagram of our approach. A sequential feature extraction algorithm process one sequence at a time whereas `FastFeatGen` can process *c* sequences at a time using *c* available cores in a computing machine. It basically distributes all genome sequences to available cores and keeps it processing in parallel which results in faster running time.

**Fig. 2:**
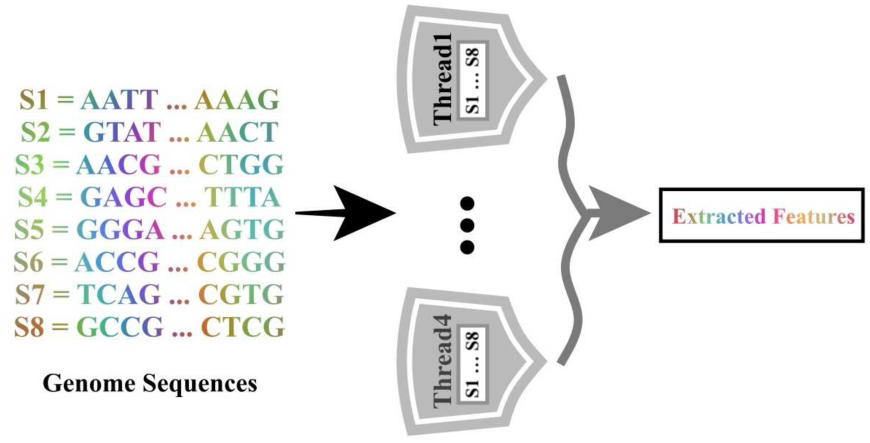
A schematic diagram of parallel feature extraction where each thread constructs a specific feature using all input sequences.

### 2.3 Machine learning algorithms

In this tool, we use several supervised machine learning algorithms either to select informative features or to train the model. For all these approaches, we incorporate the popular **sklearn** python package in our tool unless otherwise mentioned explicitly. We provide short description for each of these models below.

#### Support vector machine (SVM)

In computational genomics and proteomics, SVM is widely used for classification purpose. At first, the input dataset is transformed to high-dimensional feature space and then a kernel function maps the feature space to another dimension so that a boundary (also called margin) can separate the positive/negative classes. It creates a hyper-plane between positive and negative datasets so that margin between nearest positive samples and nearest negative samples is maximized. Nearest samples are often called support vectors and we can precisely state that the larger distances from hyper-plane equates to greater confidence levels in the predicted values. For SVM, we applied linear kernel function for feature selection and radial basis kernel function for classification purposes.

#### Random forest (RF)

RF is a popular ensemble method that is widely used for feature selection as well as for classification. Decision tree is the building block of RF which constructs rule sets over the feature space of training dataset, put class labels in the leaves of the tree, and branches denote conjunction of different rules that result in a corresponding class label. RF generally consists of a strategy to average a number of decision trees on various subsets of the dataset at training time to reduce variance and over-fitting. We allow maximum depth of RF trees to 500 in our model and use default values for other parameters.

#### Extra tree classifier (ETC)

ETC is another ensemble method which is similar to RF with few differences [12]. In ETC, each tree is trained using whole training samples instead of a subset and randomization is used while top-down splitting of a tree node i.e., a random split node is selected rather than selecting a locally optimal split node based on information gain or gini impurity. This random cut-point is selected from a uniform distribution. A final split node is selected from all randomly generated splits which achieves the maximum score. For the classification task, a prediction is made by aggregated scores of each tree by majority voting. Here, we also allow maximum depth of ETC trees to 500 to avoid possible over-fitting and default values for other parameters.

#### Neural networks (NN)

NN is one of the most popular feed-forward methods being applied in different research fields including image processing, speech recognition, bio-informatics, etc. It consists of several cascaded layers (outputs of one layer are inputs to next layer) and each layer has a finite set of nodes (neurons in human brain). Each node of a layer is connected to all nodes to its next layer which results in a fully connected network. Each connection (edge) in the network represents a weight whose optimal value is learned by an iterative optimization algorithm like stochastic gradient descent. Each node in the network adds up the products of inputs and weights and passes through the activation function, which determines how much information should proceed further to influence the predictions.

We use a variation of the NN model called deep convolutional neural networks (CNN) which is very popular in computer vision. Unlike many conventional supervised learning processes, CNN does not require manually extracted features. Rather, it can extract features by itself, which is a large advantage for automating the classification process. Manually extracted features can also be fed to CNN to build a more diversified model. For CNN, we use one-hot encoding approach to represent a sequence where ‘A’, ‘C’, ‘G’ and ‘T’ are encoded as [1, 0, 0, 0], [0, 1, 0, 0], [0, 0, 1, 0] and [0, 0, 0, 1], respectively. As a result, each 41-bp sequence is represented by a 41 × 4 matrix.

### 2.4 Feature selection

All extracted features do not contribute equally to build a better prediction model; in fact, some features do not contribute at all. We must find such irrelevant features and discard them from the feature list. We use SVM and RF for selecting and creating an important list of features that can help to train and optimize the prediction model. We use linear kernel of SVM and use a cutoff (threshold) value of 0.001 for each feature to be considered in our important feature list. Similarly, we select important features using RF based on its importance score. In the literature, RF is suggested to select a less biased or completely unbiased model [3], and many papers exist which use RF for feature selection. In our experiment, we discard any feature with zero importance from the important feature set.

### 2.5 Performance evaluation

In the literature, cross-validation is a widely used technique to build a model which reduces selection bias and overfitting problems [3]. We perform 10-fold cross-validation and jackknife testing (also known as leave-one-out cross-validation) while performing experiments for training and testing our model. In 10-fold cross-validation, the dataset is partitioned into 10 equal folds. Among these, 9 are used to train the model whereas the remaining fold is used for testing purposes. This process is repeated 10 times with a different fold selected for testing each time. In jackknife testing, *n* − 1 samples are used to train the model and the remaining sample is used for testing where *n* is the number of total samples in the dataset. This process is repeated *n* times so that each sample is considered once for testing. We use some notations of confusion matrix to define performance metrics in the following: we denote the total number of positive and negative samples by *P* and *N*, respectively; *TP*, *TN*, *FP* and *FN* represent the number of samples predicted as true positive, true negative, false positive, and false negative, respectively. We use four performance metrics, namely, Accuracy, Sensitivity, Specificity and Matthew’s Correlation Coefficient which are denoted by *Acc*, *Sn*, *Sp* and *MCC*, respectively. We express these performance metrics as following to compare our results with other tools.

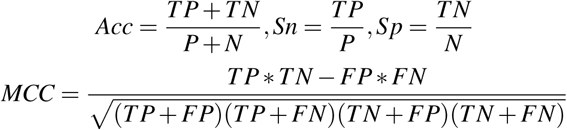

**Fig. 3:**
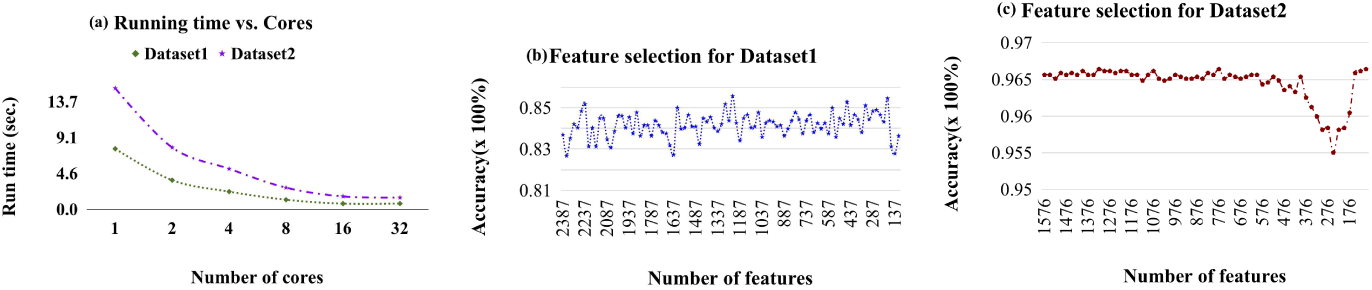
(a) Running time vs. number of cores. (b) Accuracy for different sets of features in Dataset1. (c) Accuracy for different sets of features in Dataset2.

## 3 Results and Discussion

### 3.1 Experimental setup

We perform all of our experiments in Haswell Compute Node of Cori super computer located in Berkeley lab which is configured as follows: each node has two sockets and each socket is populated with 16-core Intel®Xeon™Processor E5-2698 v3 (“Haswell”) at 2.3 GHz, 32 cores per node, 36.8 Gflops/core and 128 GB DDR4 2.13 GHz memory. We wrote the coding for our feature extraction technique in C++ and the machine learning models in Python. Our tool requires at minimum GCC version 4.9, OpenMP version 4.5, and Python version 3.6.6. We provide source code with proper documentation, results and other information in our GitHub repository.

### 3.2 Parallel feature extraction Analysis

We use shared memory parallelism for feature extraction from genome sequences which is highly scalable. To extract features from both datasets, we set parameters for NAC, PSF and DGN as 5, 30, and 28, respectively. We show the results of run time for both datasets in Fig. 3(a). For different numbers of cores, running times are reported in seconds. We observe that when we increase the number of cores, running time decreases significantly and our tool extracts features using 32 cores for the above settings within a fraction of a second. `FastFeatGen` achieves a speed-up of around 10.3x for both Dataset1 and Dataset2 using 32 threads. Our tool can be applied to a wide range of biological sequence analysis problems for feature extraction where each given dataset contains a large number of DNA/RNA/Protein sequences.

### 3.3 Feature importance analysis

As discussed in section 2.4, we select important features using linear SVM and RF. We use RF-based relevant feature selection for ETC and linear SVM-based feature selection for the SVM model. We show feature sets with accuracy for Dataset1 and Dataset2 using ETC model in Fig. 3(b) and Fig. 3(c), respectively. For Dataset1, our top-performing model contains 1237 features, among which 320 are from BPP; the rest of the features are mostly position specific. For Dataset2, our top-performing model contains 1326 features, which is higher than the case of Dataset1. 225 features are from BPP, and the rest of the features are mostly from PSF. For Dataset1, we see that *C*_26, *A*_27, and *TA*_26 are some of the features with higher importance scores. It indicates that Thymine in position 26 and Adenine in position 27 carry significant information for *N*^6^-methyladenosine sites in the rice genome. On the other hand, *G*_21 and *G*_22 are two important features for Dataset2, which indicates that Guanine in positions 21 and 22 carry significant information for the mouse genome. We observe that the accuracy of the ETC model is always higher with its combined list of features rather than its individual features. So, we use combined list of features for prediction purposes.

### 3.4 Performance analysis

#### Comparison among different learning models

To build one efficient model from different machine learning algorithms discussed in section 2.3, we conduct an extensive set of experiments. We compare SVM, ETC, NN and CNN using 10-fold cross-validation and observe that SVM and ETC models are competitive (see Tab. 1). SVM performs better than others because it uses a wide range of features, but its running time is very slow. On the other hand, ETC performs better than all other methods for a small set of features which also executes query much faster than others. The performance of NN method is comparative to SVM and ETC while CNN is the worst performer. As both datasets are small in size, CNN can not utilize its automatic feature extractions approach in depth and hence shows worse performance. We select ETC model as a representative of `FastFeatGen` for comparison with other state-of-the-art tools.

**Table 1:** Comparison among different machine learning models for Dataset1.

| Models | Accuracy | Models | Accuracy |
| --- | --- | --- | --- |
| SVM | 93.06 | NN | 80.96 |
| ETC | 84.88 | CNN | 48.41 |

#### Comparison with existing tools

We compare our model with 6mA-Pred and PseDNC, which are considered as the state-of-the-art methods for Dataset1. Following the trend of 6mA-Pred, we generate all results using jackknife test. From Table 2, we see that `FastFeatGen` (with 1237 features) achieves an accuracy of 85.56%, which is better than other existing tools. It also achieves higher Specificity and MCC which are better than other tools. Furthermore, `FastFeatGen` still outperforms existing tools in terms of Accuracy, Specificity and MCC with only 187 features.

**Table 2:** Comparison among different tools for Dataset1.

| Tools\Metrics | Accuracy | Sensitivity | Specificity | MCC |
| --- | --- | --- | --- | --- |
| FastFeatGen (1237 features) | **85.56** | 81.47 | **89.65** | **0.71** |
| FastFeatGen (187 features) | 85.45 | 81.81 | 89.09 | 0.71 |
| 6mA-Pred | 83.13 | **82.95** | 83.3 | 0.66 |
| PseDNC | 64.55 | 63.52 | 65.57 | 0.29 |

For Dataset2, we compare our method with iDNA6mA-PseKNC, which is the only tool for the mouse genome. Here, we also perform jackknife testing following the trend of iDNA6mA-PseKNC. From Table 3, we see that `FastFeatGen` is better or very competitive in all metrics.

**Table 3:**
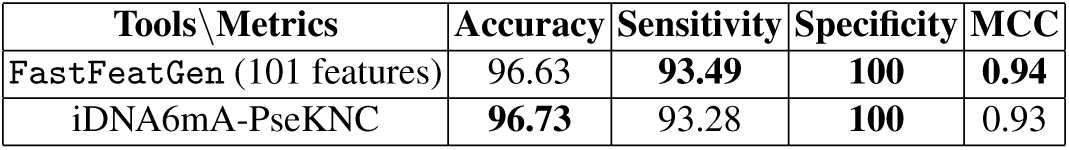
Comparison among different tools for Dataset2.

| Tools\Metrics | Accuracy | Sensitivity | Specificity | MCC |
| --- | --- | --- | --- | --- |
| FastFeatGen (101 features) | 96.63 | **93.49** | **100** | **0.94** |
| iDNA6mA-PseKNC | **96.73** | 93.28 | **100** | 0.93 |

### 3.5 Query time analysis

The general purpose of building a machine learning model is to make prediction for more unknown genome sequences, which is expected to be faster. Most of the sequence analysis tools or web-servers can not provide such facility, or authors impose restrictions on the number of query sequences. `FastFeatGen` provides a scalable solution to this problem which has no restrictions. Users can query as many sequences as they want. We again employ parallel feature extraction technique here and enable parallel job processing of ETC in **sklearn** package. In summary, `FastFeatGen` can serve 200 queries within 0.7 second(s) using 32 threads.

## 4 Conclusions

In this paper, we have introduced a novel tool called `FastFeatGen` which uses multicore processing for faster extraction of features from genome sequences. Then, it performs lucid feature selection techniques that select high quality features to feed into machine learning methods. Finally, we build a precise model using extra tree classifier which performs very well using a small set of features. We have shown that our tool performs better than state-of-the-art methods on two publicly available datasets. `FastFeatGen` achieves an accuracy of 85.56% and 96.63% for rice and mouse genomes, respectively, which are superior or competitive to current state-of-the-art methods. Our tool can predict for a wide range of new query sequences within fraction of a second which is clearly an advantage over web-server based tools.

Our future goal is to improve and apply our faster feature extraction techniques to other biological problems that involves protein or RNA sequences. We also would like to include more feature extraction techniques to `FastFeatGen`.

## Footnotes

1 https://www.ncbi.nlm.nih.gov/geo/

